# Do gametes woo? Evidence for non-random unions at fertilization

**DOI:** 10.1101/127134

**Authors:** Joseph H. Nadeau

## Abstract

A fundamental tenet of inheritance in sexually reproducing organisms such as humans and laboratory mice is that genetic variants combine randomly at fertilization, thereby ensuring a balanced and statistically predictable representation of inherited variants in each generation. This principle is encapsulated in Mendel’s First Law. But exceptions are known. With transmission ratio distortion (TRD), particular alleles are preferentially transmitted to offspring without reducing reproductive productivity. Preferential transmission usually occurs in one sex but not both and is not known to require interactions between gametes at fertilization. We recently discovered, in our work in mice and in other reports in the literature, instances where any of 12 mutant genes bias fertilization, with either too many or too few heterozygotes and too few homozygotes, depending on the mutant gene and on dietary conditions. Although such deviations are usually attributed to embryonic lethality of the under-represented genotypes, the evidence is more consistent with genetically-determined preferences for specific combinations of egg and sperm at fertilization that results in genotype bias without embryo loss. These genes and diets could bias fertilization in at least three not mutually exclusive ways. They could trigger a reversal in the order of meiotic divisions during oogenesis so that the genetics of fertilizing sperm elicits preferential chromatid segregation, thereby dictating which allele remains in the egg versus the 2^nd^ polar body. Bias could also result from genetic- and diet-induced anomalies in polyamine metabolism on which function of haploid gametes normally depends. Finally, secreted and cell-surface factors in female reproductive organs could control access of sperm to eggs based on their genetic content. This unexpected discovery of genetically-biased fertilization in mice could yield insights about the molecular and cellular interactions between sperm and egg at fertilization, with implications for our understanding of inheritance, reproduction, population genetics, and medical genetics.

---

Our understanding of inheritance in sexually reproducing organisms assumes, with good evidence, that the combination of egg and sperm at fertilization is largely independent of their genetic content. This equal transmission of alternative alleles through meiosis in heterozygotes ensures a balanced parental genetic contribution to offspring at each generation. Mendel’s First Law captures this principle, which is one of the few that applies generally in biology. Independent segregation and random union of gametes at fertilization are foundations of classical, quantitative, population, evolutionary and medical genetics.^1-3^

The most prominent exceptions to random segregation are the rare naturally-occurring examples of transmission ratio distortion (TRD) that have been described in fungi,^4^ corn,^5^ flies,^6-10^ mice,^11-16^ humans,^17, 18^ and other species. ^19-22^ Biased sex ratios have also been reported.^23-29^ These exceptions arise despite strong selective pressures that strive to maintain normal segregation^2^ and sex ratios^30, 31^ Based on haploid effects in gametes, one allele is preferentially transmitted to offspring at the expense of other alleles. TRD is usually the property of one sex, driving allelic preference regardless of the genetics of the mating partner. Reproductive performance is not reduced because the normal number of gametes is produced. TRD may arise during gene and chromosome segregation in meiosis (meiotic drive), gametogenesis (gamete competition), or embryonic development (preferential lethality). These examples of TRD probe the nature of Mendel’s First Law by illuminating genetic, molecular and cellular mechanisms that underlie meiosis, recombination, gametogenesis, and early development.^1-3, 13, 32-36^ Many of these ‘selfish genetic’ systems are composed of several closely linked elements that not only lead to preferential transmission of the chromosome on which they are carried, but also confer sterility or lethality on homozygous carriers, thereby preventing fixation at the cost of reduced population fitness.^11^ Our listing of TRD examples must be limited to these systems because otherwise the driver would quickly replace their wildtype (WT) alleles in the population, evidence of their competitive advantage would be lost, and we would be ignorant of their history.

With spontaneous, induced and engineered single gene mutations, one of the first tasks is to assess consequences on viability, fertility and other phenotypes.^37-39^ Absence of mutant homozygotes is usually accepted as evidence for induced lethality and reduced numbers of heterozygotes is taken as evidence for a detrimental dosage effect (Fig. 1). But sometimes the fit to Mendelian segregation is not explicitly examined. Litter size, which should be reduced proportionately to the number of missing genotypes, is often not reported. Backcrosses, which can provide information about parent-of-origin effects on gametogenesis and embryogenesis, are sometimes not included in study designs. As a result, whether particular cases of non-Mendelian segregation result from lethality or from other phenomena such as TRD remains ambiguous, despite claims in publications.

**Figure 1.**
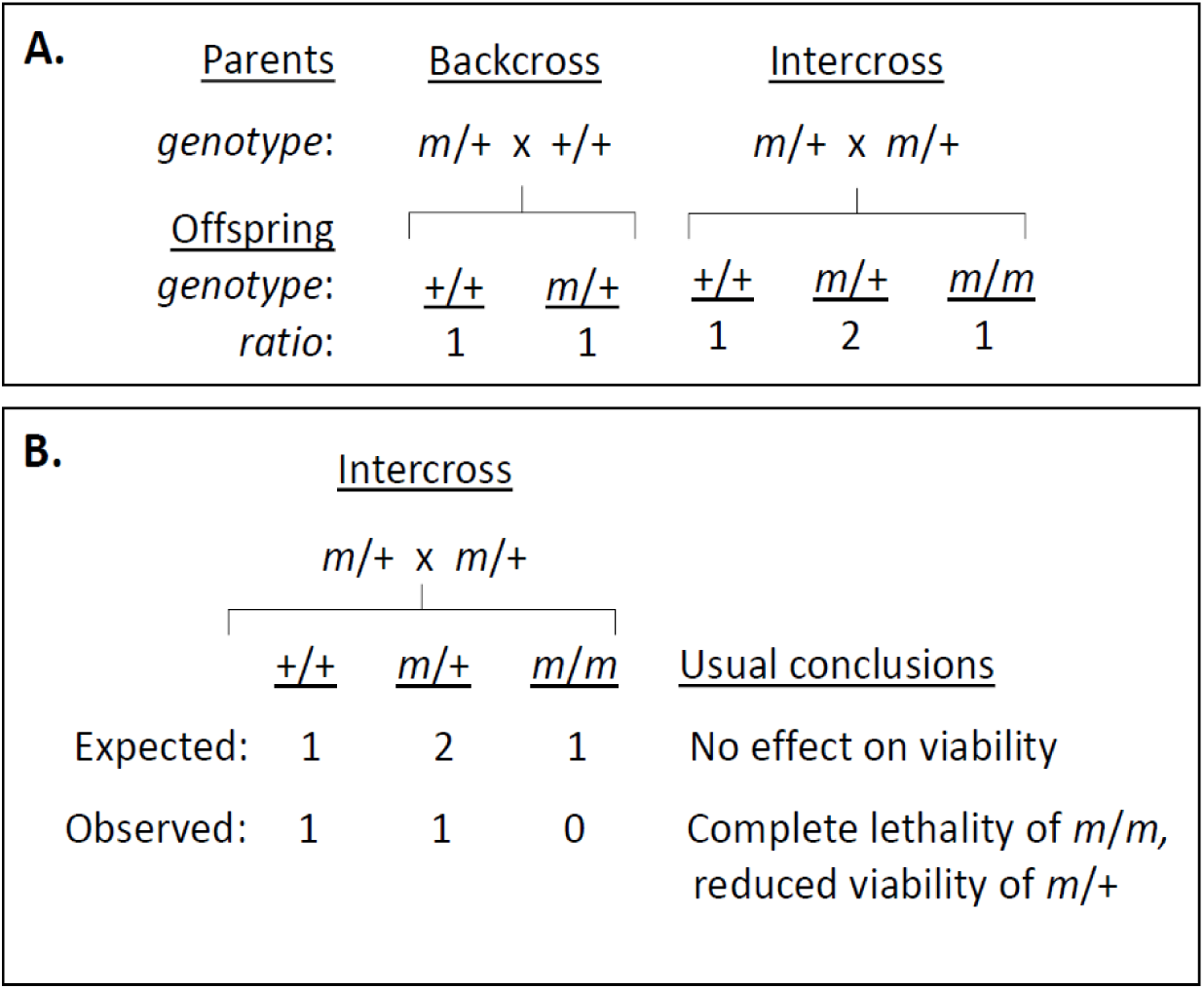
Regular and irregular outcomes of Mendelian segregation. A. Conventional segregation in backcrosses and intercrosses for a single gene with two alleles, wildtype (+) and mutant (*m*). B. Irregular segregation in an intercross with loss of all *m*/*m* homozygotes and half of +/*m* heterozygotes.

Consider an early controversy in mammalian genetics. Cuenot studying absence of pure yellow segregants in crosses between mice heterozygous for the dominant yellow (*A^y^*) mutation argued that gametes carrying the yellow allele never join at fertilization.^40^ By contrast, Castle and Little, based on considerations of both segregation ratios and litter size, correctly concluded that homozygous yellow mice fail to complete development, with reduced litter size providing the critical evidence for lethality.^40, see also 41^ As Castle and Little showed, a full analysis is needed to establish with confidence the basis for unusual segregation.

Before the introduction of molecular techniques for genotyping sperm and eggs, Mendel’s Law could only be tested indirectly by genotyping the next generation, after gametes become zygotes. Implicit and untested assumptions are sometimes made that genotypic ratios among offspring correctly reflect meiotic products among gametes. Random union of gametes at fertilization is one of these assumptions. Many life stages and events such as gametogenesis, fertilization, embryogenesis and post-natal development occur in the interim between mating and offspring surveys, any of which could lead to departures from expectations. Indeed, when Castle-Little considerations are applied to relevant data for new mutants, evidence consistent with Cuenot’s hypothesis of biased fertilization is sometimes found.

Our work on epigenetic inheritance in mice and a selective review of the mouse literature revealed strong evidence for TRD based on the genetic constitution of both egg and sperm at fertilization. Briefly, depending on mutations in any of 12 genes (Box 1), either too many or too few heterozygotes were found among intercross progeny, together with absent or deficient mutant homozygotes, without evidence for dead embryos or reduced litter size (Tables 1, 2). Normal segregation in backcrosses between heterozygotes and WT homozygotes argues that meiosis and gametogenesis function normally in each sex. Six cases involve single spontaneous or engineered mutations on an inbred genetic background. In another case, biased segregation was found only in crosses involving a pair of mutant genes (epistasis). Finally, five cases involved dietary folic acid supplementation of mice carrying single-gene mutations affecting neural tube development, where segregation was biased on one diet but normal on the alternative diet, with similar litter sizes and rates of prenatal lethality. These unusual results suggest that fertilization is genetically biased towards particular gamete combinations. Here, evidence for TRD resulting from non-random union of gametes in mice is reviewed, and then possible mechanisms and genetic implications are considered.

**Table 1.** Summary of genetic effects. Average transmission ratio (TR) for backcrosses (bc’s) and intercrosses (ic’s), calculated as the ratio of mutant heterozygotes (*m*/+) to wildtype (+/+) with a Mendelian expectation of 1 (=1:1) for backcrosses and 2 (=2:1) for intercrosses (expected ratio in parentheses). Effect size assesses the difference between the average TR and Mendelian expectations. Cohen’s *w* for a goodness-of-fit test was used as a standardized measure of effect size, with a difference between observation and expectation of *w* = 0.10 as the threshold for declaring a weak effect, 0.30 for a medium effect and 0.50 for a strong effect,^44^ with medium and strong differences highlighted in gray. Additional information is provided in Suppl. Table 1.

| Genes | Observed TR (*m/+*: +/+) |  | Average effect size |  | Litter size (no. litters) |  | Conclusion for intercrosses |
| --- | --- | --- | --- | --- | --- | --- | --- |
|  | backcross | intercross | backcross | intercross | backcross | intercross |  |
| *Ago2* | 0.89 | 1.03 | 0.07 | 0.34 | 4.9 (216) | 3.9 (33) | too-few heterozygotes |
| *Dnd1* | 0.89 | 1.35 | 0.04 | 0.20 | 5.7 (48) | 5.2 (19) |  |
| *A1cf* | 1.10 | 6.39 | 0.05 | 0.35 | 5.8 (178) | 5.9 (51) | too-many heterozygotes |
| *Ddx1* | 1.15 | 8.15 | 0.07 | 0.48 | 8.1 (81) | 7.3 (52) |  |
| *Ppp2cb* | 0.79 | 3.48 | 0.12 | 0.23 | 7.6 (73) | na |  |
| *Pum1* | 1.68 | 5.17 | 0.25 | 0.36 | 6.9 (26) | 6.9 (53) |  |

## What is the evidence for fertilization bias?

### Genetics

Two kinds of TRD were found, one with a deficiency (too-few), the other an excess (too-many) of heterozygotes. Central to this evidence is the expectation that departures from Mendelian segregation that result from embryo loss should reduce reproductive performance as measured by the number of offspring produced, here ‘litter size’ because evidence is from mice (Fig. 1). For example, lethality of all *m/m* homozygotes and half of the *m/+* heterozygotes should reduce litter size by half, that is 1:1:0 rather than 1:2:1. Dead embryos should also be found. Biased genotype distributions together with normal litter sizes and absence of dead embryos argue for non-random fertilization rather than lethality. As expected, most genetic variants segregate normally and show reduced litter size in proportion to genotype bias, both in our hands and in the literature (informatics.jax.org; www.komp.org), suggesting that biased fertilization is exceptional and results from specific rather than general dysfunctions. All of the evidence reported here involves single gene mutations on an inbred strain background. Two cases of fertilization bias have been reported previously,^42, 43^ although the responsible genetic factor is not known in either case.

To summarize evidence for non-random fertilization, emphasis was placed first on testing departures from Mendelian expectations, namely 1:1 in backcrosses and 1:2:1 in intercrosses (Fig. 1), and then on measures of the nature and magnitude of these departures (Table 1 for genetic effects and Table 2 for gene-folic acid interactions, see also Suppl. Tables 1 and 2 for complete data, analytical methods, and results). Given the absence of mutant homozygotes in many intercrosses (Suppl. Table 1), transmission ratio (TR) was based on the relative number of heterozygous (*m/+*) to homozygous wildtype (+/+) offspring with an expected ratio of 2 (2:1) for intercrosses and 1 (1:1) for backcrosses. Effect size is an important but often neglected measure of phenotypic differences and was used here to provide a normalized quantitative measure of departures from expectations for genetic and gene-diet effects.^44^ According to accepted standards,^44^ effects sizes >0.10 are classified as ‘small’, >0.30 ‘medium’, and >0.50 ‘strong’. These measures are independent of sample size.

### Single genes, simple cases

In these cases, 1:1 and 1:2:1 segregation is expected for backcrosses and intercrosses respectively if inheritance is Mendelian. In addition, litter sizes will be reduced if departures from expectations results from embryonic lethality, but will be normal if fertilization is biased. For *Dnd1* and *Ago2* mutants, segregation was highly unusual with significant deficiencies of *m/+* and *m/m* genotypes among intercross progeny – too-few heterozygotes (Table 1, see also Suppl. Table 1). For backcrosses, TRs were close to 1:1 expectations and effect sizes were small, whereas for intercrosses TRs were closer to 1 than 2 and effect sizes were medium (Table 1). Heterozygotes that are missing in intercrosses are found in expected numbers in backcrosses. Embryonic lethality does not account for the observed genotype ratios because litter sizes were similar among intercrosses and backcrosses (Suppl. Table 1). For *Dnd1*, neither mutant homozygotes nor dead embryos were found at embryonic day E3.5.^45^ For *Ago2*, a 25% reduction in litter size is consistent with the reported lethality of *m/m* mutant homozygotes,^46^ but not with the observed 50% deficiency of *m*/+ heterozygotes.

For *A1cf*, *Ppp2cb* and *Pum1*, highly significant excesses of *m/+* (too-many) heterozygotes were found together with absence of mutant homozygotes among intercross progeny (Table 1, see also Suppl. Table 1). For backcrosses, TR was close to 1:1 expectations and effect sizes were small, whereas for intercrosses, TR ranged from 3.1 to 9.7, instead of the expected ratio of 2, and effect sizes were large (Table 1). Despite these differences, average litter size was remarkably similar for backcrosses and intercrosses (Suppl. Table 1). For *Pum1*, *m/m* embryos were not detected at E3.5 and litter size did not differ between intercrosses and backcrosses.^47^ For *A1cf*, the heterozygote excess ranges from 3- to 5-fold, instead of the expected value of 2, based on two reports;^48, 49^ Loss of homozygous embryos between E3.5 - 4.5 does not account for excess heterozygosity or for normal litter size.^48^ Although litter size was not reported for *Ppp2cb,^50^* the highly significant heterozygote excess (>3:1) in intercrosses versus backcrosses is striking and consistent with results for *A1cf*, *Ddx1* (see below) and *Pum1.* Normal segregation in backcrosses with mutant heterozygotes shows that gametes are produced in comparable (1:1) numbers and functionality in each sex.

Excess heterozygosity is a curious and unexpected observation. A possible explanation involves lethality of wildtype and mutant homozygotes, but normal litter size argues against this. The question then is whether there are circumstances under which TRD results in too-many heterozygotes. Three models were considered (Fig. 2): (1) TRD in one sex but not the other, (2) identical TRD in both sexes, and (3) opposing TRD (similar magnitudes but opposite directions in the two sexes). The one-sex scenario is included only for reference; it does not apply to the present circumstances, otherwise TRD would have been found in both intercrosses and backcrosses. Using various TRs for the WT allele, we calculated the genotypic ratios and then, to facilitate comparison with observed genotype distributions (Tables 1,2), calculated TR, the ratio of heterozygotes to WT. In intercrosses without TRD, this ratio should be 2 (= 2 heterozygotes to 1 WT). Results for one sex- and both sex-TRD are remarkably similar with an excess of heterozygotes arising when TR <0.50, and a deficiency when TR>0.50 (Fig. 2). Thus, excess heterozygosity is found when the wildtype allele is favored and a deficiency when the *m* allele is favored. For *Dnd1* and *Ago2*, too-few heterozygotes arose only when the WT allele is favored. The more interesting opposing-TRD case identified two conditions under which too-many heterozygotes might arise, namely when the wildtype allele is favored in one sex and disfavored in the other, that is TR<0.5 or TR>0.5. Opposing TRD has been reported, usually as a result of ‘sexual antagonism where the advantage of a mutant allele differs, being favored in one sex and disfavored in the other.^51-55^ Interestingly, several pathways and tissues that are involved in sexual antagonism are also implicated in fertilization bias.^56, cf. Box 1^ These models highlight circumstances under which the observed deviations from expectations might arise and provide a guide to interpreting mechanistic studies.

**Figure 2.**
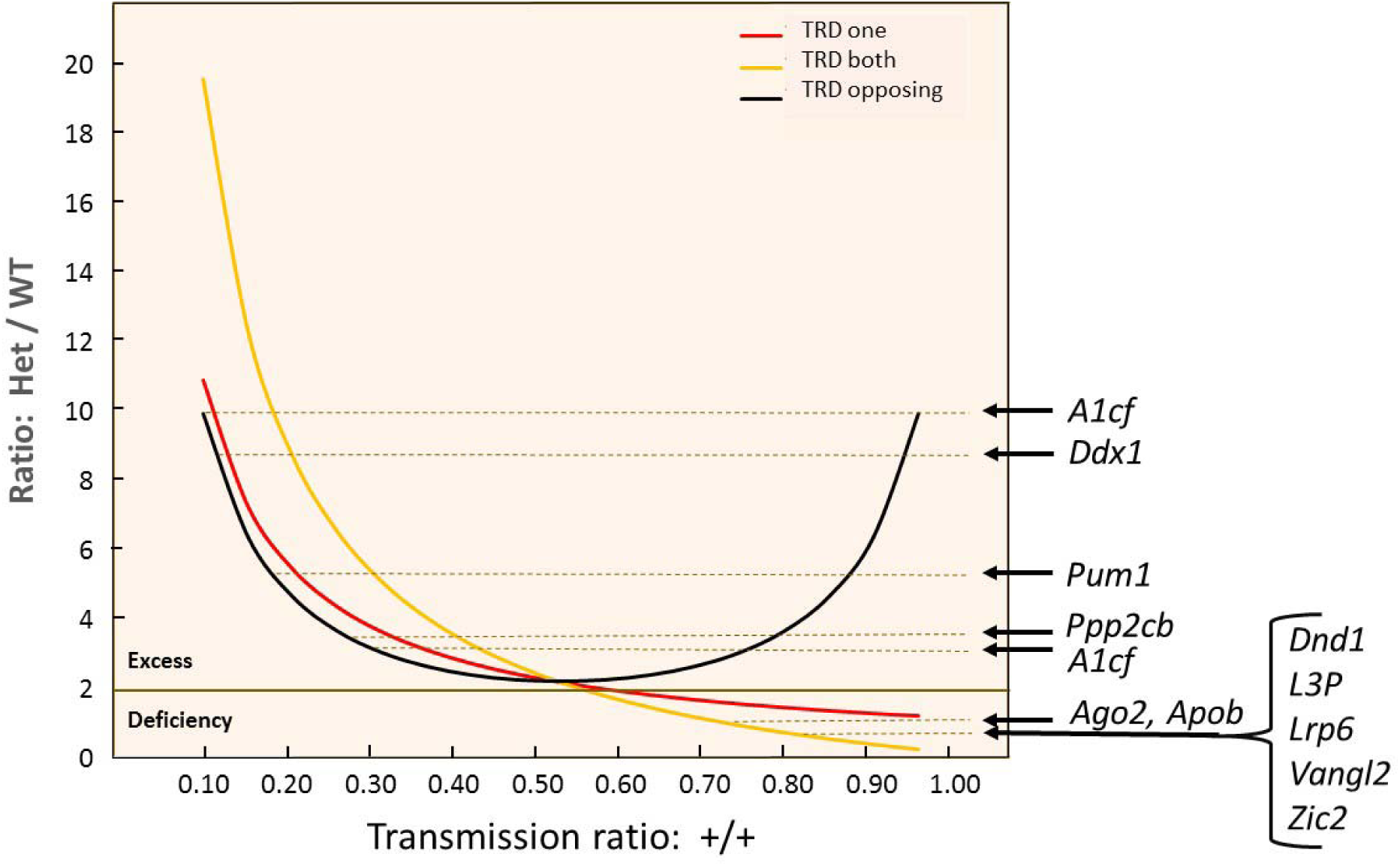
Origins of ‘too-many’ and ‘too-few’ heterozygotes. Three models (TRD one-sex, TRD both-sexes, TRD opposing) were examined to determine the scenarios under which ‘too-few’ and ‘too-many’ heterozygotes might arise. Horizontal line at “2” denotes the transmission ratio for 2 mutant heterozygotes to 1 wildtype homozygote expected in intercrosses without TRD. Dashed lines show the observed ratio of heterozygotes to wildtype for each gene (see Tables 1,2 and also Suppl. Tables 1,2) *A1cf* is shown twice because evidence is available from two publications.^48, 49^ Methods and interpretations are provided in the text.

### Ddx1 - complicated single gene effect

An engineered deficiency of DEAD box 1 helicase (*Ddx1*) and an induced epigenetic change in its WT allele provide strong evidence for biased fertilization.^57, 58^ For the engineered mutation, *m/m* embryos were missing with no homozygotes detected at E3.5 in test crosses. In addition, a substantial deficit of WT segregants relative to *m/+* heterozygotes was also observed in some crosses (Table 3, suppl. Table S1), leading the authors to conclude that the engineered mutation induced a modified, perhaps paramutated,^58^ allele (designated ‘*’) that leads to lethality in both WT (+/+) and */* embryos resulting from crosses involving a */+ parent. However, review of litter sizes for test and control crosses revealed remarkably little evidence for embryo loss that would account for the putative loss of *m/m*, */* and +/+ embryos. Strongly biased transmission without embryo loss argues that preferential fertilization is a more likely explanation.

**Table 2.** Summary of gene-folic acid effects. Two diets (2 ppm and 10 ppm, parts per million) were tested on a panel of single gene mutations that cause neural tube defects (NTDs). Five genes provided evidence for biased fertilization. Expectations (Exp) are based on Mendelian segregation ratios for intercrosses. Exp 2 corresponds to 2 mutant heterozygotes to 1 wildtype homozygote and Exp 1 to 1 mutant homozygote to 1 wildtype homozygote. Effect size is Cohen’s *w* for a goodness-of-fit test^44^ (see Table 1 for details). Additional information is provided in Suppl. Table 2.

| Genes | Folic acid dose | TR |  | Effect size | Litter size (no. litters) | Diet with effect |
| --- | --- | --- | --- | --- | --- | --- |
|  |  | m/+ : +/+ | m/m : +/+ |  |  |  |
| *L3p* | 2 ppm | 1.43 | 0.39 | 0.30 | 7.2 (13) | 2 ppm |
|  | 10 ppm | 2.00 | 1.14 | 0.06 | 4.2 (10) |  |
| *Zic2* | 2 ppm | 1.56 | 0.57 | 0.20 | 6.1 (18) |  |
|  | 10 ppm | 2.57 | 0.95 | 0.14 | 5.7 (10) |  |
| *Apob* | 2 ppm | 2.00 | 1.13 | 0.05 | 6.2 (16) | 10 ppm |
|  | 10 ppm | 0.95 | 0.68 | 0.33 | 5.8 (18) |  |
| *Lrp6* | 2 ppm | 1.97 | 0.54 | 0.22 | 7.9 (20) |  |
|  | 10 ppm | 1.56 | 0.29 | 0.36 | 8.5 (22) |  |
| *Vangl2* | 2 ppm | 3.15 | 1.15 | 0.16 | 4.3 (16) |  |
|  | 10 ppm | 1.39 | 0.48 | 0.26 | 4.4 (15) |  |

**Table 3.**
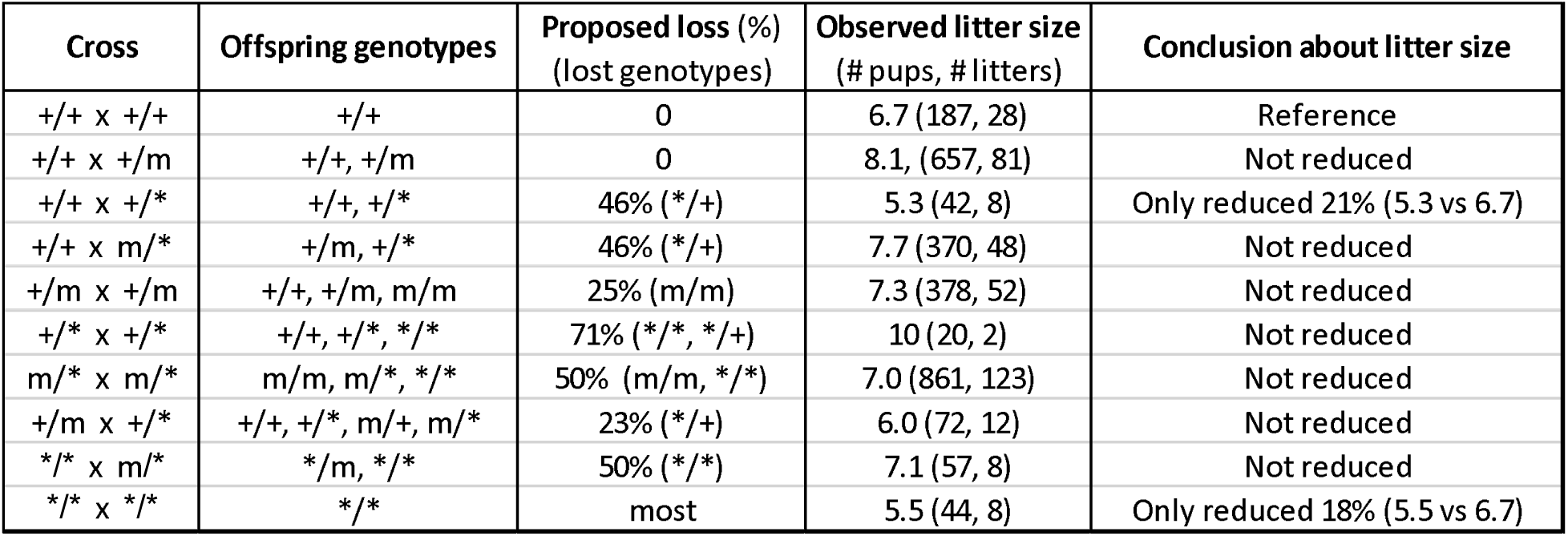
*Ddx1* segregation. Three alleles are shown, wildtype (+), mutant (*m*) and modified wildtype (*). Various crosses were used to examine the effect of *m* and * on embryonic viability. Offspring genotypes are shown for each cross. Proposed loss is based on the hypothesis that the following genotypes would result in embryonic lethality: *m*/*m* and */* and where m/*, m/+ and */+ are viable.^57^ Loss is summarized as the deviation (percentage) from Mendelian expectations for these genotypes in each cross. ‘Conclusion about litter size’ is based on comparing the observed litter size with the ’reference’ for the wildtype cross. Data are from Hildebrandt et al.^57^ and from R. Godbout (pers comm).

**Table 4.**
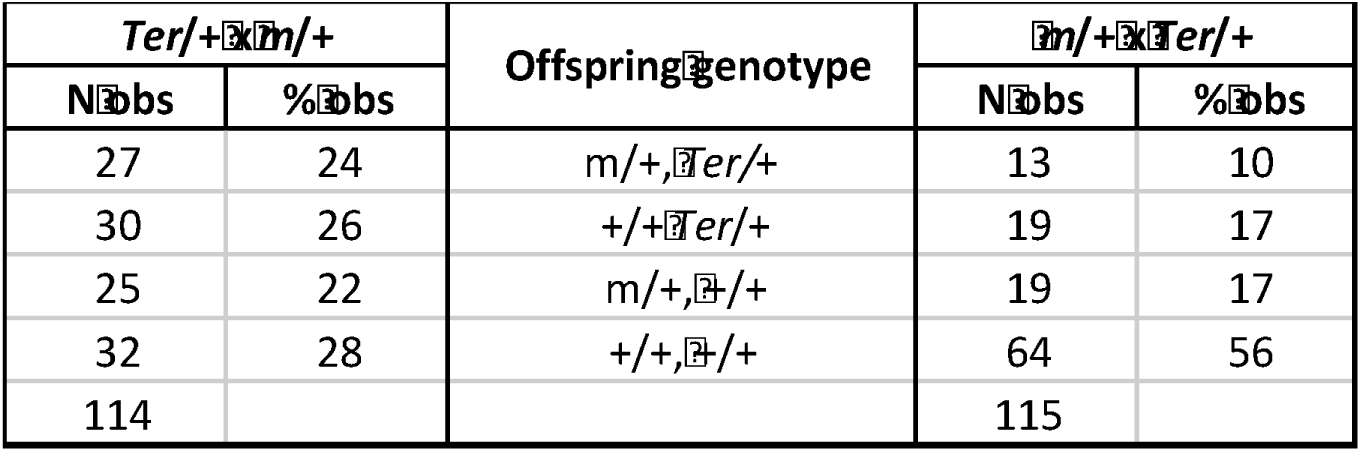
Segregation in Apobec1, Dnd1^Ter^ intercrosses. *Ter* is a spontaneous mutation in the *Dnd1* gene.^61^ Reciprocal crosses were made between *Apobec1* targeted deficiency mutation (*m*) and *Dnd1*^Ter^. Crosses are ‘female’ x male’. N obs is number observed for each genotype and cross. % obs is the percentage for that genotype among all offspring for each cross. For two segregating genes in these intercrosses a 1:1:1:1 Mendelian ratio is expected. Results are from Nelson et al.^59^

### *Apobec1 and Dnd1* - a complicated two-gene effect

The last genetic example emerged in tests to determine whether *Apobec1* and *Dnd1* interact to modulate inherited risk for spontaneous testicular germ cell tumors (TGCTs; see below for additional information about TGCT origins, genetics and biology).^59^ These genes independently affect risk in a conventional and an epigenetic manner.^59^ But whether they interact in the genetic sense was uncertain. Surveys for TGCTs among intercross offspring revealed unexpected evidence for biased fertilization (Table 4). With maternal heterozygosity for *Dnd1* and paternal heterozygosity for *Apobec1*, offspring occurred in the expected (1:1:1:1) Mendelian ratio for two independently segregating genes. But results for the reciprocal cross revealed a marked deficiency of all single- and double-mutant genotypes. If the number of WT (+/+, n=64) progeny is accepted as the proper reference for the three other genotypic classes, where 1:1:1:1 = 64:64:64:64, then only 27% (51/192, where 192 = 3 x 64) of the expected single- and double-mutant segregants was found. Again, litter size did not differ between the reciprocal crosses.^59^ Remarkably, both heterozygous and homozygous mutants for each gene are fully viable in separate crosses (suppl. Table S1) ^45, 48, 60, 61^. Thus, in this but not the reciprocal cross, the majority of mutant segregants is missing, with evidence for full viability in other crosses and with no evidence for reduced litter size.

### Gene – folate diet interactions

A related class of TRD involves dietary effects on single gene models of neural tube defects (NTDs). The beneficial effect of dietary folate supplementation on a common birth defect is one of the greatest achievements in epidemiology.^62^ NTDs such as anencephaly and spina bifida are the second most common congenital defect, occurring in ∼1 per 1000 live births and often leading to disability and mortality.^63, 64^ Mothers and fetuses frequently show reduced folate and elevated homocysteine levels.^62, 65^ Dietary folate supplementation before and during pregnancy significantly reduces the occurrence and severity of cases, but many (∼50%) remain resistant to the beneficial effects of folate supplementation.^62, 64, 66^ Reliable prediction of individual response to supplementation is currently impossible.

Several studies examined the effects of dietary supplementation with folic acid on development of the neural tube in mouse models.^67-69^ Some respond favorably to folic acid,^70^ several respond to other nutrients such as methionine and inositol,^71, 72^ but as in humans many are resistant.^67-69, 73^ In the study design for these studies, developmental response to various nutrients is tested among intercross progeny. In some cases, +/*m* heterozygotes are maintained on a test or control dietary formulation, bred together (+/*m* x +/*m*), and developmental consequences among +/+, +/*m* and *m*/*m* offspring evaluated, where offspring are expected to occur in a 1:2:1 Mendelian ratio. In other cases, pregnant +*/m* females are treated only during critical developmental windows and then embryos or offspring examined. The former is the only protocol that is instructive about possible dietary effects on gametogenesis and fertilization. Attributing effects to diet simply involves comparing results for the high versus low dosage treatment groups on the same defined genetic background. Efficacy is indicated if the number (proportion) of affected *m*/*m* individuals is reduced.

During studies to identify mouse NTD models that are resistant to the benefits of folic acid treatments, two examples of fertilization bias were found.^74^ Reanalysis of published reports then revealed three additional NTD models that show biased fertilization in response to folic acid treatment (Table 2). In several cases (*Apob*, *Lrp6*, *Vangl2*), significant deviations are found in the high folic acid (10 ppm; parts per million) test, but in other cases (*L3P*, *Zic2*) deviations were found in the low (2 ppm) test, with similar litter sizes for each mutant on the two diet protocols (suppl. Table S2). TRs approximated expectations for genes showing normal segregation and effect sizes were small, whereas for the genes showing departures from expectations, TRs were more divergent and effect sizes were medium (Table 2, see also Suppl. Table 2). None showed too-many heterozygotes. Departures with dietary supplementation were not as strong as the genetic results (Table 1), perhaps because optimal supplementation levels were not tested and perhaps because dietary consumption and metabolism differ among the various cohorts and are generally more difficult to control.

These results raise a provocative question, namely does folate correct a developmental defect in the neural tube, or do other explanations apply such as reducing the incidence of cases by biasing fertilization away from at-risk genotypes?^74^ Interestingly, for most models, the percentage of affected *m/m* was similar in the test and control protocols, suggesting that supplemental folic acid did not reduce the proportion of affecyed mutant homozygotes.^75, 76^ In humans where NTD genetics is not as clearly understood as in mouse models, with genetic heterogeneity a significant issue and isolated cases common, distinguishing a protective effect versus biased fertilization would be difficult.

### Gene functions

Six of the seven TGCT genes encode proteins that are directly involved in various aspects of RNA biology: *A1cf* – RNA editing,^77, 78^ *Apobec1* – RNA editing,^77^ *Ago2* – RNAi,^79^ *Dnd1* – miRNA control^80^ *Ddx1* – RNA helicase,^57^ and *Pum1* – translation repression^47^ (Box 1). The seventh gene, *Ppp2cb*, is a serine/threonine phosphatase^50^ (Box 1). Four show inherited epigenetic effects on TGCT risk (*A1cf, Apobec1*, *Ago2*, *Dnd1*) and one shows epigenetic effects on embryonic viability (*Ddx1*). Neither *Ppp2cb* nor *Pum1* have been tested for TGCT or epigenetic effects. These proteins have specific RNA targets that are in turn effectors of developmental and physiological functions. These functions could be shared or distinct in males and females depending on the nature of the targeted RNAs in each sex. Identifying these targets is critical for understanding the ways that these genes control gamete choice at fertilization.

By contrast, the five NTD-genes appear to have diverse, seemingly unrelated functions with no obvious theme (*Apob, L3P, Lrp6, Vangl2, Zic2*; Box 1). Perhaps RNA genes and *Ppp2cb* are directly involved in molecular and cellular mechanisms of gametogenesis and fertilization, whereas the various NTD genes sensitize folate and homocysteine metabolism to adverse interactions with pathways that directly affect gamete interactions at fertilization.

### Competition between diploid and haploid phases

Discoveries about biased fertilization are relevant to theories concerning sexual antagonism, namely the contrasting priorities between diploid organisms and their haploid gametes. Diploids strive for reproductive success versus other diploids, whereas haploid gametes compete with each other for fertilization success. Because gametic competition could reduce parental fertility, diploid cells seek to reduce functional differences among gametes by limiting their transcription and translation. As long as gametes are functionally equivalent, diploids have the advantage over their haploid gametes. It is noteworthy then that many of the genes that bias fertilization affect aspects of RNA biology that impact translation (Box 1). Partial loss of function in mutant heterozygotes might enable phenotypic differences among gametes, leading to gamete competition and biased fertilization.

Functional effects confined to haploid gametes are critical for The four products of meiosis produced in males usually have an equal chance of fertilization, in part because syncytia provide small intercellular channels through which developing sperm share molecules, thereby minimizing impact of haploid effects.^81^ However, gametes carrying a t-haplotype, which are a classic example of TRD in the mouse,^11, 13^ produce two critical elements that control sperm motility in a haploid-specific manner. Gametes that carry a *t*-haplotype produce molecules that pass through syncytial bridges to hyperactivate Rho signaling in both *t*- and +-bearing sperm, thereby compromising their motility. However, *t*-bearing sperm also produce a protective, haploid-specific variant of SMOK1 (sperm motility kinase) that does not pass through bridges and protects *t*-bearing but not +-bearing sperm, thereby providing a motility advantage to *t*-bearing sperm.^35, 82, 83^ Because such effects are intrinsic to heterozygous males and would be found in both backcrosses and intercrosses, a corresponding effect would need to operate in females to be a sufficient explanation for biased fertilization.

### Centromere- or gene-specific effects

Sometimes TRD results from preferential segregation of centromeric elements that guide chromosome movement and segregation during karyokinesis.^9, 84-86^ Genes that are closely linked to the centromere would also show TRD with the degree of distortion depending on the recombination distance from the centromere and with genes located 50 cM or more showing normal 1:1 segregation. However, several genes that are located far from the centromere show substantial TRD and overall there is little evidence that TRD declines as a strong function of recombination distance from the centromere (Fig. 3). In addition, dispersed chromosomal locations for these genes (Tables 1,2) argue against a selefish gene complex, as is found frequently with other TRD systems.^13, 34^ Together these observations are consistent with gene-specific TRD rather than hitch-hiking resulting from close linkage to centromere-driven elements.

**Figure 3.**
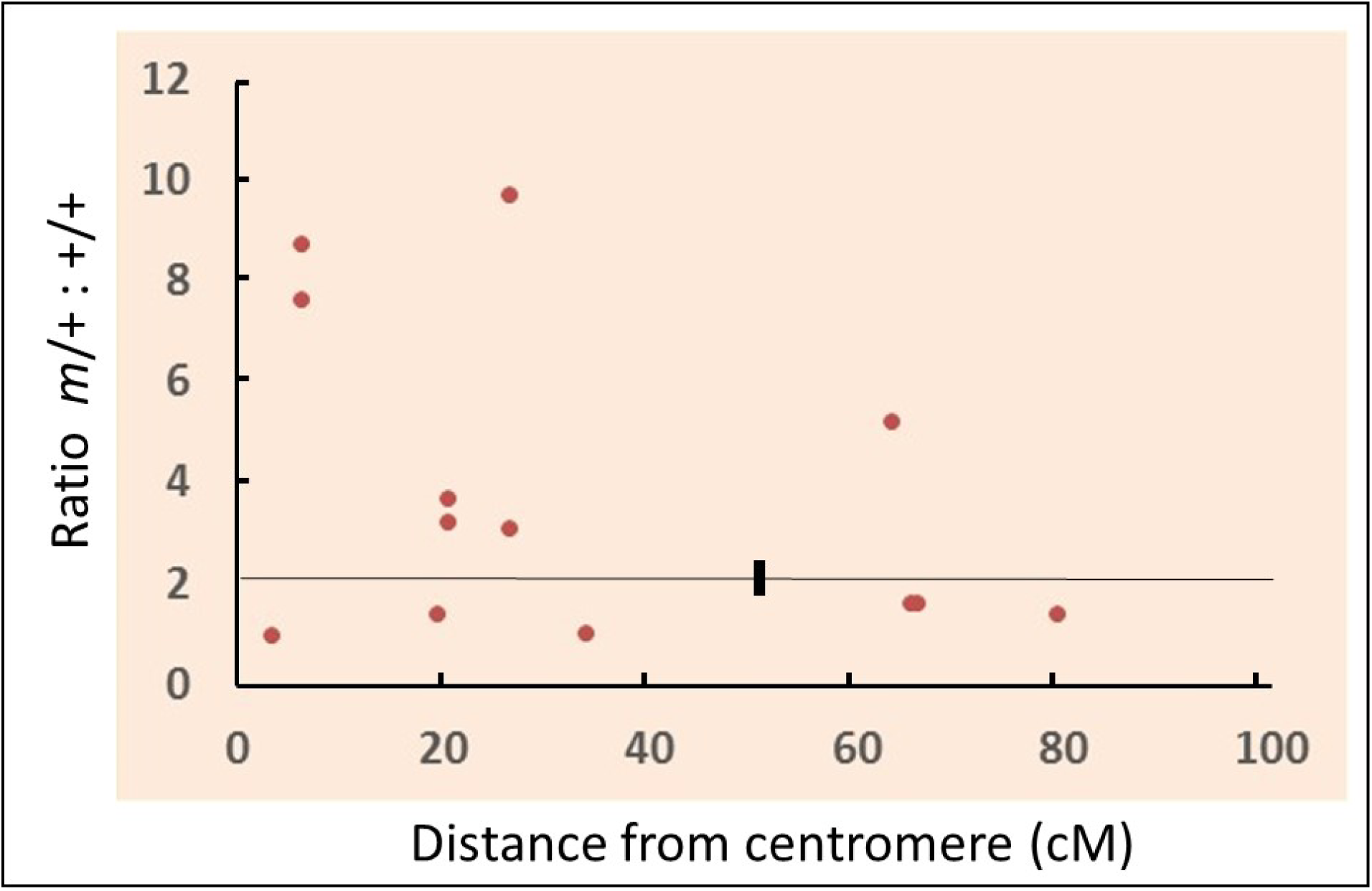
Relation between transmission ratio (*m*/+ : +/+) and distance from centromere. Transmission ratios are from Tables 1 and 2; recombination distances are from the Mouse Genome Database (informatics.jax.org). The horizontal line at 2 (= 2:1) corresponds to the expected ratio of *m*/+ heterozygotes to +/+ homozygotes expected with Mendelian segregation. A black box marks 50 cM from the centromere. The correlation was modest (r = -0.37) between the transmission ratio and gene distance from the centromere, accounting for only 11% of the variation and suggesting that centromeric drivers, if any, had modest effect on transmission ratios. In addition, the four genes beyond the 50 cM mark are unlikely to result from any centromeric effect.

### Litter size

Although many factors such as the number of fertilized eggs, pre- and postimplantation mortality, and uterine capacity can affect litter size, various evidence shows that the primary determinant is the number of ovulated eggs. Selection for larger litter size increases ovulation rate,^87-90^ while selection for increased ovulation rate results in larger litters.^91^ Inbreeding reduces litter size because of fewer ovulated eggs.^92^ Unilateral ovariectomy reduces litter size by 50%,^93^ arguing against eggs held in reserve to compensate for failed fertilizations and embryo loss. In parallel, eggs mature within ovarian follicles that rupture to release eggs at ovulation, with the number of growing follicles determined by host genetics long before and independent of fertilization. Thus the number of ovulated eggs available for fertilization appears to be the primary determinant of litter size.

### Lethality

Three lines of evidence argue against lethality as the explanation for departures from Mendelian expectations (Fig 1). First, litter size is not reduced in intercrosses versus backcrosses for any genes that bias fertilization (Tables 1-3). Again, if mutant homozygotes are embryonic lethal, then litter size should be reduced 25%, and if half the heterozygotes are also missing, then litter size should be further reduced to 50%. Second, in some cases (*Dnd1* ^45^ , *Ddx1* ^57^, *Pum1* ^47^), surveys at E3.5 failed to find mutant homozygotes or dead embryos. Third, loss of particular genotypes is not based on their inherently deleterious nature, suggesting that negative genotypic selection is not involved. Mutant heterozygotes that are missing among intercross progeny are found in expected numbers among backcross progeny (Table 1), in reciprocal crosses (Table 4), or on alternative folic acid diets (Table 2).

### What are the mechanisms?

#### Reversed meiosis

Normally, meiotic divisions in females are ordered the way we have been taught, first the reductional division (MI), then the equational division (MII) (Fig 4). During ovulation, primary oocytes resume meiosis and over the next several hours complete MI and arrest at metaphase in MII.^94^ Homologous (non-sister) chromatid pairs segregate at MI with one product going to the secondary oocyte and the other to the first polar body, which may divide again at MII.^95, 96^ Fertilization triggers completion of MII with sister chromatids segregating to the second polar body and the oocyte.^96^

**Figure 4.**
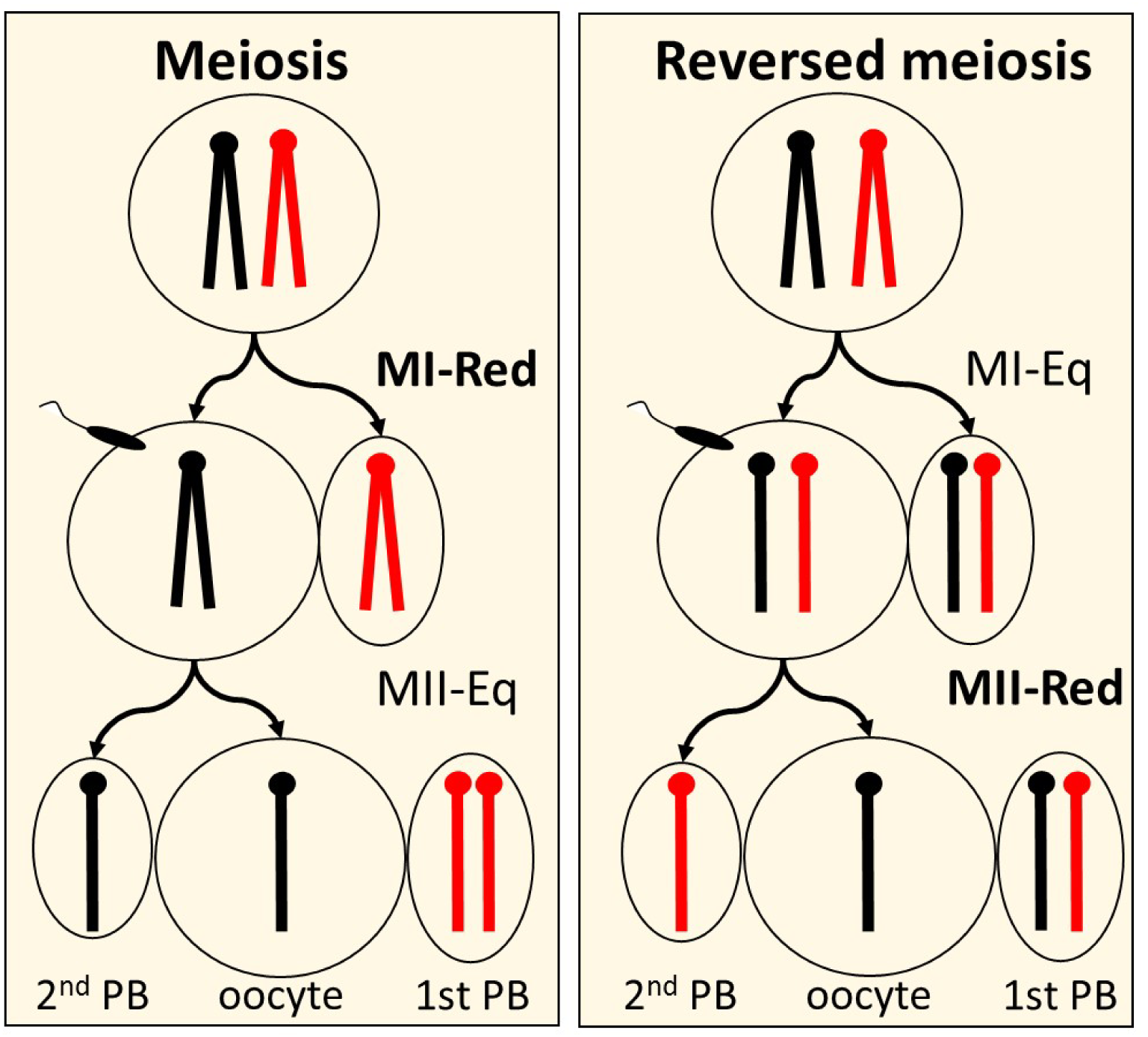
Regular and reversed meiosis. Alternative chromatids are marked in red and black, MI-reductional (MI-Red) precedes MII-equational (MI-Eq) in regular meiosis, whereas in reversed meiosis MI-equational occurs first.^95, 96^ Recombination is not included in these scenarios, although normally ∼40% of chromatids at the haploid phase have a crossover.^209^ MII arrests at metaphase until sperm entry at fertilization. PB – polar body.

Recently, ‘reversed meiosis’ was reported after induced ovulation in humans.^97^ Reversing the order of meiotic divisions during oogenesis, with the equational division occurring at MI rather than MII, leaves the secondary oocyte heterozygous for marker genes at fertilization (Fig 2). The genetics of fertilizing sperm could then bias MII segregation with one chromosome preferentially remaining in the oocyte and the other segregating to the second polar body. Although formal tests have yet not been reported in mice, the Agulnik study is consistent with reversed meiosis.^42^

Interestingly, genes such as *Ago2* and *Ppp2cb* that control chromosome segregation in mitosis are strong candidates for determining the sequence of meiotic divisions.^98-100^ The alternative hypothesis that meiosis is conventional cannot account for fertilization bias because completion of MI prior to fertilization makes the reductional division independent of the mating partner and the genetic content of fertilizing sperm. Bias should therefore be evident in both backcrosses and intercrosses. Whether meiotic reversal serves an adaptive purpose or is an anomaly resulting triggered by reduced gene dosage and physiological stresses such as hormonal treatments to induce ovulation is unclear.

The length of the haploid phase in spermatogenesis and oogenesis places conditions on the mechanisms that might contribute to biased fertilization. In mammals, the haploid phase is long in males, with the MI division arrested during embryonic development and recommencing at puberty. Spermatogenesis then continues throughout reproductive life. The haploid phase begins with completion of meiosis in the testis and continues as they mature and capacitate and while they pass through the epididymis, vas deferens, urethra and uterus to fertilization in the oviduct, a period that can last several days. By contrast, the haploid phase is remarkably short in females, lasting only from completion of MII, which fertilization triggers, until female and male pronuclei fuse. This brief window raises the likelihood that fertilization drives the bias in oocytes.

#### Polyamines

Polyamine metabolism connects dietary and molecular effects on folate metabolism with functional consequences in haploid gametes. Most of this evidence involves spermatogenesis; the evidence for oogenesis is meager by comparison. Polyamines (PAs) such as spermine, spermidine, putrescine and cadaverine are low molecular weight organic molecules that have at least two amino groups. They are present in all cells and most are associated with nucleic acids.^101^ They are involved in transcription, translation, histone modifications, autophagy, apoptosis and many other molecular, cellular and physiological functions.^102-106^ Given their interdependent roles in essential molecular, epigenetic and cellular functions, the polyamine and folate pathways are highly conserved and highly regulated, from yeast to plants and mammals, with sometimes overlooked roles in gametogenesis and fertility.^104, 107-112^ The substrates for polyamine synthesis are ornithine, which is derived from arginine and proline, and S-adenosylmethione (SAM), which is part of the homocysteine cycle in folate metabolism (Fig 5). SAM is best known as the methyl donor for all methylation reactions for nucleic acids (DNA, RNAs, including tRNAs), proteins (including histones), lipids and other molecules.^101, 113-115^ Moreover, by utilizing acetyl CoA, polyamine catabolism magnifies the effects of folate deficiency (Fig 5). Acetyl CoA is the substrate for synthesizing betaine, which is the alternative methyl donor to synthesize SAM from homocysteine. However, despite the essential role for methylation in many molecular and cellular functions, cells preserve polyamine metabolism at the expense of methylation, at least under *in vitro* and *in vivo* stress conditions.^116-120^

**Figure 5.**
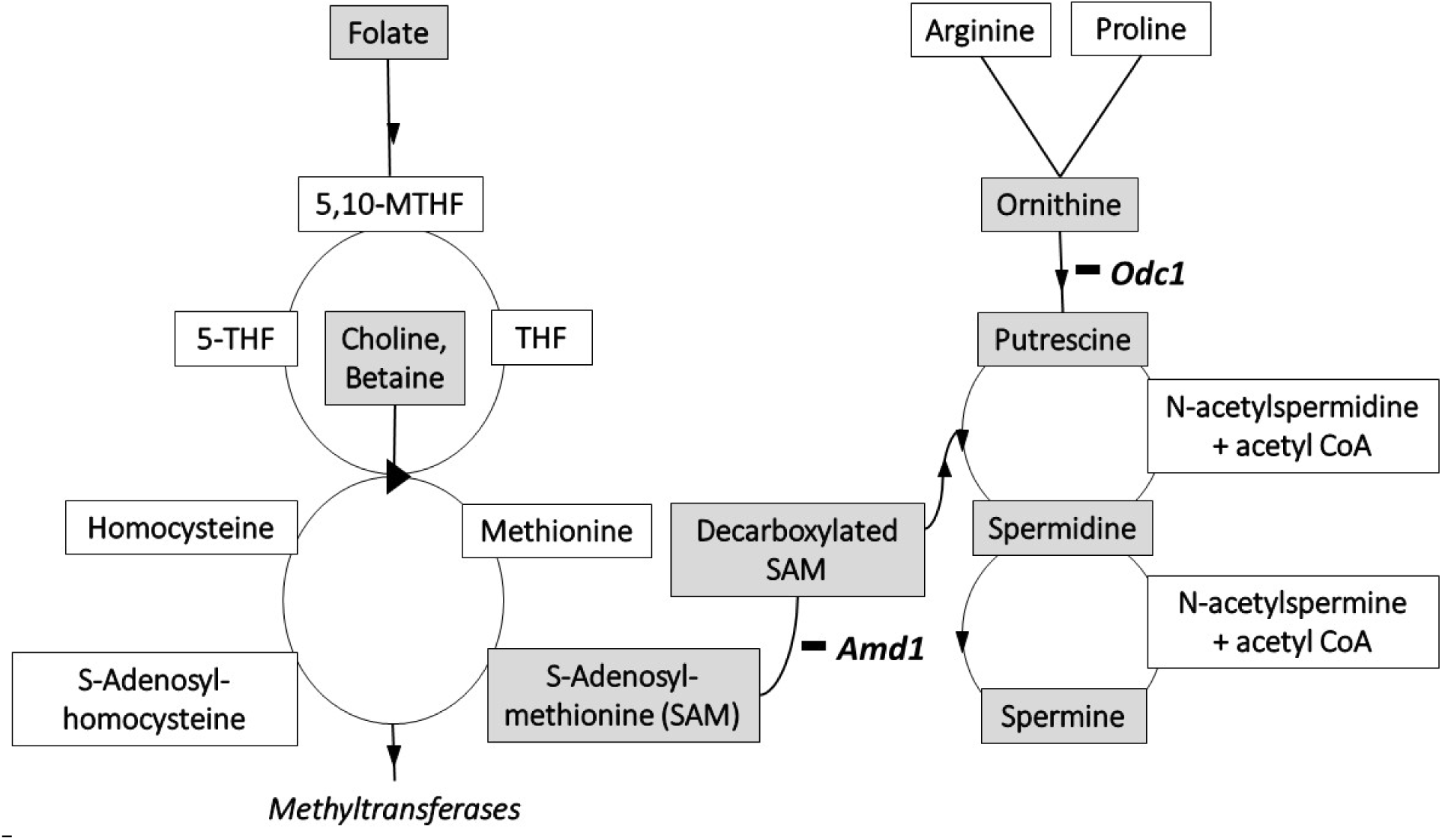
Folate and polyamine metabolic pathways. (from ref ^65^). Gray highlights key molecules. Abbreviations: 5,10-THF (5,10-methylenetetrahydrofolate), 5-THF (5-tetrahydrofolate), THF (tetrahydrofolate). Folate is the primary methyl donor; choline and betaine are alternative methyl donors. *Amd1* (S-adenosylmethione decarboxylase 1) links folate-homocysteine metabolism with polyamine metabolism by converting S-adenosylmethionine to decarboxylated S-adenosylmethionine, which with putrescine produces spermidine. *Odc1* (ornithine decarboxylase 1) converts ornithine to putrescine and is the heavily regulated rate-limiting step in polyamine synthesis. Catabolism of spermine and spermidine consumes acetyl CoA, which also serves as a substrate for choline and betaine synthesis.

Functionality of haploid gametes depends heavily on polyamine metabolism in many species. In Arabidopsis, the MAT3 S-adenosylmethionine synthase gene is expressed in pollen, with mutants showing reduced pollen tube growth and seed set as well as changes in polyamine biosynthesis, tRNA levels, and histone methylation.^112^ In humans, polyamine deficiency results in infertility, which can be corrected with SAM or polyamine supplementation.^104, 121-123^ In cattle, spermine is essential for acrosomal function.^122^ Over-expression of ornithine decarboxylase 1 (ODC1), the rate-limiting step in polyamine synthesis, also leads to infertility.^124-127^ Contrary to dogma, sperm translate nuclear-encoded mRNAs by using mitochondrial-type ribosomes; blocking translation reduces sperm motility, capacitation, and *in vitro* fertilization (IVF) rate.^128^ Several polyamine genes are expressed in haploid gametes.^104^ ODC antizyme 3 (OAZ3) is a testis-specific inhibitor of ODC1^124, 126, 127, 129, 130^ and deficiency leads to sperm that cannot fertilize.^126, 127^ Antizyme inhibitor 2 (AZIN2) blocks the inhibitory effects of both OAZ3 in haploid cells^131^ as well as ODC1 over-expression.^132^ SAT1 (spermidine/spermine N1- acetyltransferase) and OAZ1 are differentially expressed with folate supplementation in the LRP6 mouse NTD model^74^ and OAZ1 is differentially expressed in sperm from folate-deficient mice.^133^ Finally, the primary inputs for polyamine metabolism, namely the enzyme S-adenosylmethionine decarboxylase 1 (AMD1), which catalyzes the conversion of SAM to decarboxylated SAM, and the amino acids arginine, ornithine and proline are critical for pluripotency control in embryonic stem cells (ESCs) (Fig. 5).^134-140^ Given the close lineage relations between germ cells and ESCs,^141, 142^ similar effects in gametes would not be surprising. Polyamine metabolism is therefore a strong candidate for biased fertilization given its established impact on haploid gametes and its strong connection to folate metabolism.

#### Sperm in the epididymis, gametes in the oviduct

The issue here is whether reproductive organs and sperm-egg recognition provide opportunities for gametic selection based on haploid effects^143^ Semen as well as oviductal and uterine fluids contain various proteins, nucleic acids, and small molecules that provide a chemically appropriate environment for sperm maturation and capacitation for fertilization and for implantation of genetically non-self embryos.^93, 144-151^ Because mating often occurs prior to ovulation, sperm can be in the oviduct hours before ovulation, often needing just minutes from ejaculation to arrive in the oviduct.^152^ Sperm and eggs induce gene expression changes in the oviduct that alters the biochemistry of oviductal fluids^153^ and that in turn restrict sperm access.^143, 154^ Millions of sperm are released at ejaculation but usually less than 100 reach the oviduct.^155^ Not only do sperm compete for access to eggs,^156, 157^ the oviduct can select sperm based in part on their genetic content, including sex chromosome (X or Y)^154, 158^ and on chromatin stability.^159^

Signaling before contact between sperm and egg has obvious advantages for haploid gametes. There are reports that small transiently expressed peptides attract a minority of sperm that are only briefly responsive, presumably representing the 10% of sperm that are appropriately capacitated.^160, 161^ Thermotaxis,^162^ chemotaxis,^143, 154^, and signaling molecules including olfactory receptors,^163^ trace-amine-associated receptors (TAARs),^164, 165^ cannabinoid receptors,^166^ and calcium receptors^167 168, 169^ have been reported in gametes and gonads. These presumably play a role in sperm-egg attraction, but the evidence is limited both about mechanisms and especially about the impact of genetic variation on molecular interactions.

In Drosophila and other species, success of fertilizing sperm depends on the genetics of both the male and female mating partners, suggesting that ligand-receptor interactions are critical.^170, 171^ But how genetic variation affects affinity and signaling in these ligand-receptor pairs is largely unknown. Once sperm penetrate the glycoprotein coat surrounding the egg, sperm and egg must recognize each other and fuse membranes. The zona pellucida hardens irreversibly to prevent polyspermy.^172^ Presumably fertilization bias must occur before membrane fusion and zona pellucida hardening, otherwise the conceptus would either persist as a viable embryo, or be lost with a corresponding reduction in litter size. Recognition between sperm and egg is based on ligand-receptor binding between Izumo1 (sperm) and Juno (egg), absence of either protein leads to infertility.^173-175^ Although Juno, which is a member of the folate receptor family, no longer binds folate,^174^ residual functions might still depend on folate levels. Unexplained anomalies between other ligand-receptor pairs such as loss of one but not the other member of the pair resulting in infertility suggests that models of sperm-egg recognition remain incomplete.^176^ In at least one instance, union of sperm and egg brings together two distinct proteins that act together as a dimer to suppress mutagenesis in early embryos.^177^

Several approaches have been employed to define mechanisms for sperm-egg recognition. For example, ENU mutagenesis has been used to find genes controlling gametogenesis and fertilization. ^178-180^ Although many of these mutated genes affect germ cell biology and meiosis, genes affecting fertilization were not found.^181^ Fortuitously, our genetic studies of epigenetic TGCTs risk and NTD dietary response discovered several such genes, thereby enabling new genetic approaches for studying mechanisms of gamete function at fertilization.

### Epigenetics in the germline

Biased fertilization may be a previously unrecognized manifestation of genes that control TGCT susceptibility, epigenetic inheritance, and related germline abnormalities. The first evidence for fertilization bias was discovered during work on the control of inherited TGCT risk in the 129 family of inbred strains.^59, 182, 183^ These TGCTs are models for several classes of TGCTs in humans,^142, 184^ and, like gametes, originate from the germ cell lineage during fetal development.^185, 186^ Their common origin suggests that they share similar vulnerabilities to perturbations. Perhaps no other lineage undergoes such dramatic transitions in developmental potential and with such profound implications for health, fertility, and perpetuation of the germline across generations.^187^ The germline is unipotent until fertilization when it transitions to pluripotent embryonic cells that in turn differentiate into specialized somatic and germ cell lineages. ^188-190^ Various mechanisms preserve genomic integrity and developmental capacity of the ‘mother of all stem cell lineages’^191^ by repairing DNA defects,^192^ maintaining cellular conditions,^193^ suppressing transposon activity,^194-199^ and programming epigenetic state.^200, 201^ Failure of pluripotency control can lead to precocious differentiation of germ cells,^202, 203^ spontaneous transformation of germ cells during fetal development,^142, 185, 186, 204^, infertility,^49, 205, 206^ and other reproductive disorders.^204^ Dysfunction can also lead to inherited epigenetic changes that affect risk for TGCTs and urogenital abnormalities in offspring and later generations in the absence of genes that originally triggered these transgenerational effects.^49, 59, 182, 183^ Genetic anomalies together with dietary, hormonal and other environmental influences may induce germline dysfunctions that manifest as conventional genetic effects, unconventional epigenetic inheritance, and now preference for particular gamete combinations at fertilization.^49, 59, 74^

## Perspectives

Given this evidence, how might biased fertilization work? We can identify essential elements and conditions, but we can be less certain about the ways that these genes affect fertilization.

Two components are needed for bias, one in females, the other in males; neither alone is sufficient. Each element predisposes gametes to bias, but bias arises only when predisposed gametes come together at fertilization.

In females, mutations in any of these twelve genes, acting alone (*A1cf, Ago2, Dnd1^ko^, Ddx1, Ppp2cb, Pum1*), with each other (*Apobec1, Dnd1^Ter^*), or in combination with particular diets (*Apob, Lrp6*, L3P, Zic2, Vangl2), could reverse the sequence of meiotic divisions with the equational preceding the reductional division instead of the usual order.^97^ This would leave the secondary oocyte heterozygous, so that the genetics of fertilizing sperm could then ‘drive’ one allele (chromosome) preferentially to the polar body while retaining the other allele (chromosome) in oocyte. The reversed MII division, and hence which allele is retained in the egg, would be resolved only after fertilization, and thus may not be independent of the genetics of fertilizing sperm. Reversed meiosis is critical, otherwise biased fertilization would occur in females regardless of the genetics of their mating partner. The source of sperm is also critical, with fertilization bias found in sperm from mutant heterozygotes but not wildtype homozygotes. It is unclear whether specific gene functions control the order of meiotic divisions or whether their mutations, sometimes including diet effects, induce epigenetic changes that result in reversed meiosis.

Special conditions must also apply in males. Unlike meiotic drive and gamete competition, fertilization bias cannot simply result from preferential chromosome segregation or from gamete dysfunction or deficiency. Otherwise TRD would be found in both backcrosses and intercrosses and would be independent of the genetics of the female mating partner. Instead. backcross evidence (Table 1, suppl. Table 1) and diet tests (Table 2, suppl. Table 2) show that both mutant and wildtype sperm genotypes have an equal chance of fertilizing eggs from wildtype females, demonstrating that these sperm do not have intrinsic deficits that compromise their functionality. Instead, bias is found only when these sperm encounter eggs in heterozygous females. Moreover, interactions between predisposed eggs and sperm must occur prior to sperm entry, otherwise the fate of the egg is fixed with the hardening of the zona pellucida to prevent polyspermy. The chemical environment surrounding sperm and eggs could contribute to preferred fertilization. However, this environment would need to differ between heterozygous and homozygous females. Perhaps signaling between combinations of ligands and their receptors mediate these interactions. These molecules are only recently beginning to be characterized.^207^ Whether genetic variation in these proteins, especially in the ligand-binding site,^208^ remains to be examined.

The twelve genes are evidence of causality, with functions ranging from RNA-mediated gene silencing, RNA editing, mRNA deadenylation and microRNA regulation to lipid transport, cilia function and signal transduction (Box 1). Mutations, including simple dosage effects (hemizygosity), in any of these 12 genes, either alone (Table 1), together (Table 4) or with gene-diet interactions (Table 2), are sufficient to bias fertilization away from gamete combinations that would preserve Mendelian expectations. However these genetic effects must occur in both mating partners to bias fertilization; either alone results in Mendelian transmission. This suggests that heterozygosity for these mutations or exposure to particular diets predisposes gametes to bias. But this bias is only realized when predisposed sperm and eggs are present together at fertilization.

The ways that these mutations affect gametes at fertilization is unclear. Because many of these genes have multiple targets, it is possible that hemizygosity compromises the same targets and functions in both sexes, or that they target different but functionally relevant genes in each sex. This interpretation is based on the specific functions and targets of each gene, gene pair, and gene-diet combination. An alternative model is based on induced epigenetic changes in the germline. Considerable evidence shows that hemizygosity for many of these genes results not only in increased risk for germ cell tumors, but also for parent-of-origin and transgenerational effects on tumor risk in offspring and later generations. Perhaps hemizygosity alone, rather than specific gene functions, induces widespread epigenetic changes in the germline, resulting not only in germ cell tumors but also in epigenetically distinct gametes that together bias fertilization. The role of folate metabolism in DNA methylation and polyamine metabolism in histone modifications is consistent with this epigenetic interpretation.

Four steps are needed to transfer genetic and epigenetic information from one generation to the next through the germline. Meiosis converts the chromosome complement from diploid to haploid in the parental generation. Gametogenesis provides a cellular vehicle for the haploid genome. From a pair of haploid gametes, fertilization restores diploidy in the zygote. Finally, the germline is set aside early in embryonic development to renew these steps in the offspring generation. TRD has been reported for three of these steps, - meiotic drive, gamete competition, and preferential embryo survival. In each case, dysfunction in one sex is sufficient for TRD that is largely independent of the genetics of mating partners. The evidence reviewed here provides examples for TRD in the third step – fertilization, where genetic variants, acting in both sexes and in some cases depending on environmental (dietary) conditions, control the combination of gametes that join at fertilization to create zygotes. Historically, the genetics of fertilization has been largely resistant to molecular studies.^207^ Discovery of genes and gene-diet conditions that bias fertilization may be a breakthrough in understanding mechanisms of sperm-egg interactions at fertilization.

We are familiar with the elaborate, elegant and sometimes extravagant rituals that organisms often use to attract mates and assess fitness. Perhaps gametes woo too.

## Acknowledgements

I thank Rosaline Godbout and her group for generously sharing litter size and segregation data from Hildebrandt al al.^57^ and for sharing her thoughts about their study and the interpretation proposed here. I also thank David Crews, Mary Ann Handel, and especially Hamish Spencer and Monika Ward for comments on a draft of this paper. Andy Clark, Aimee Dudley, John Eppig, Fernando Pardo de Villena, Jasper Rine, Jesse Riordan, Carmen Sapienza and Patrick Stover made helpful suggestion and insights. NIH grants NICHD Pioneer Award <u>DP1 HD075624</u>, NCI CA75056 and NINDS NS058979 supported this work.

---

**Supplemental Table 1.**
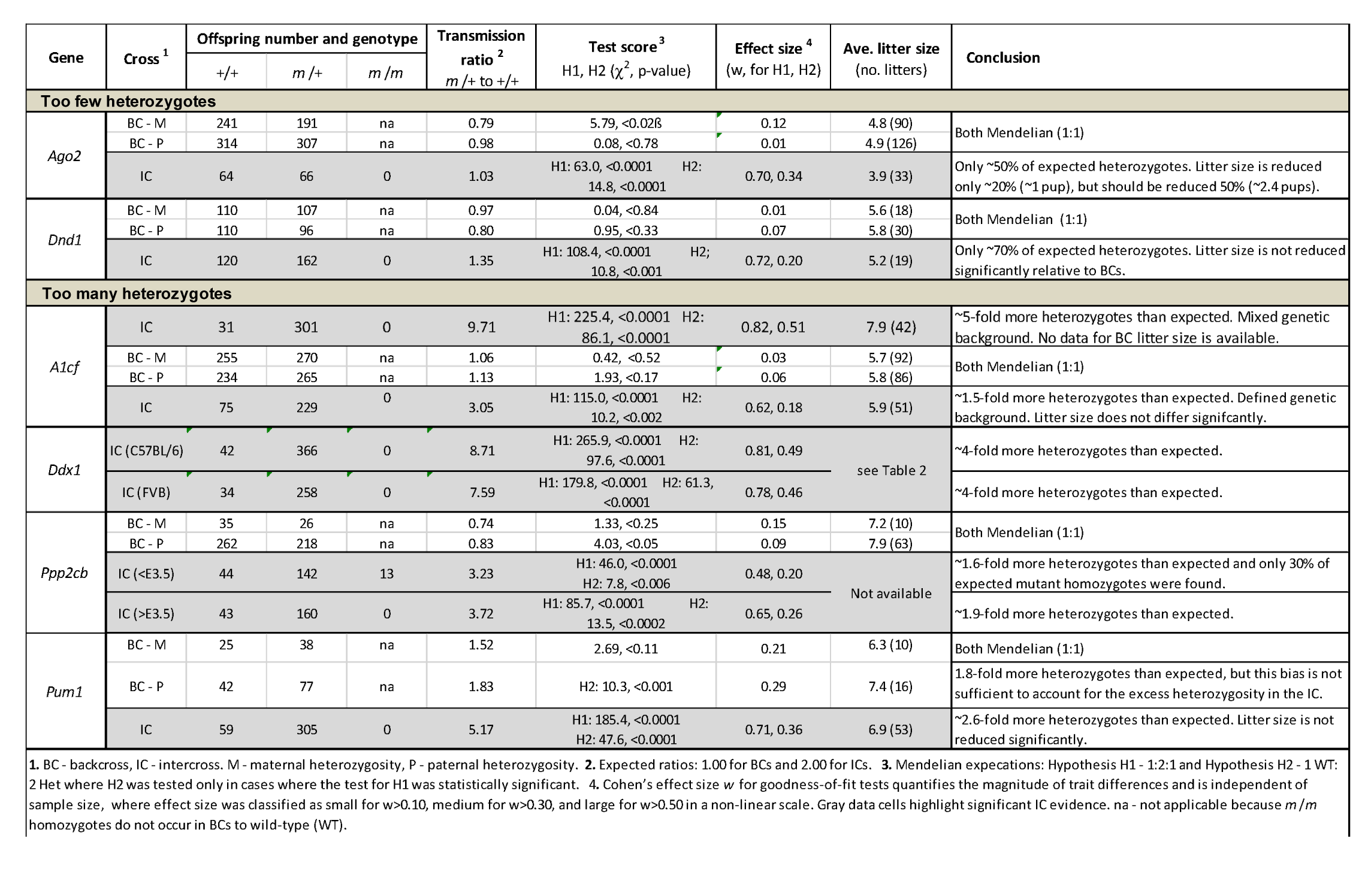
Genetic effects on segregation.

**Supplemental Table 2.** **Gene-diet effects on segregation.** Data are derived from Nakouzi et al.,^74^ Gray et al.,^75^ and Marean et al.^76^

| Gene | Folic acid | Offspring number and genotype |  |  | TR<sup>1</sup><br>*m* /+ to +/+,<br>*m* /*m* to +/+ | Test score<sup>2</sup><br>H1, H2 (χ<sup>2</sup>, p-value) | Effect size<sup>3</sup> (w) | Ave. litter size<br>(no. litters) | Expected<br>litter size<sup>4</sup> | Conclusion |
| --- | --- | --- | --- | --- | --- | --- | --- | --- | --- | --- |
|  |  | +/+ | *m* /+ | *m* /*m* |  |  |  |  |  |  |
| 2 ppm folic acid effect |  |  |  |  |  |  |  |  |  |  |
| *L3P* | 2 ppm | 28 | 40 | 11 | 1.43, 0.39 | H1: 7.3, <0.03<br>H2: ns | 0.30 | 7.2 (13) | 2.6 (0.61) | Significant deficiencies of *m* /+ and *m* /*m*, but with increased litter size. |
|  | 10 ppm | 14 | 28 | 16 | 2.00, 1.14 | ns | 0.06 | 4.2 (10) |  |  |
| *Zic2* | 2 ppm | 56 | 84 | 32 | 1.56, 0.57 | H1: 6.8, <0.04<br>H2: ns | 0.20 | 6.1 (28) | 3.9 (0.69) | Significant deficiencies of *m* /+ and *m* /*m*, but with similar litter size. |
|  | 10 ppm | 38 | 98 | 36 | 2.57, 0.95 | ns | 0.14 | 5.7 (30) |  |  |
| 10 ppm folic acid effect |  |  |  |  |  |  |  |  |  |  |
| *Apob* | 2 ppm | 24 | 48 | 27 | 2.00, 1.13 | ns | 0.05 | 6.2 (16) |  | Significant deficiencies of *m* /+ and *m* /*m*, but with similar litter size. |
|  | 10 ppm | 40 | 38 | 27 | 0.95, 0.66 | H1: 11.1, <0.004<br>H2: 11.3, <0.0008 | 0.33, 0.33 | 5.8 (18) | 3.3 (0.54) |  |
| *Lrp6* | 2 ppm | 37 | 73 | 20 | 1.97, 0.54 | ns | 0.22 | 7.9 (20) |  | Significant deficiencies of *m* /+ and *m* /*m*, but with similar litter size. |
|  | 10 ppm | 55 | 86 | 16 | 1.56, 0.29 | H1: 20.8, <0.0001<br>H2: ns | 0.36 | 8.5 (22) | 4.9 (0.62) |  |
| *Vangl2* | 2 ppm | 13 | 41 | 15 | 3.15, 1.15 | ns | 0.16 | 4.3 (16) |  | Significant deficiencies of *m* /+ and *m* /*m*, but with similar litter size. |
|  | 10 ppm | 23 | 32 | 11 | 1.39, 0.48 | H1: ns; H2: ns | 0.26 | 4.4 (15) | 2.7 (0.62) |  |
| 1. **Transmission ratio** (TR) of genotypes to assess departures from Mendelian expectations for *m* /+ x *m* /+ intercrosses, where offspring genotype ratios *m* /+ to +/+ should be 2:1 and *m* /*m* to +/+ should be 1:1. 2. See Suppl. Table 1. 3. See Suppl. Table 1. 4. The following calculation was used to estimate the expected litter size if lethality accounted for the departures from expectations. First, assume the number of wildtype (+/+) offspring is the proper number in each test group and therefore is one of the '1' terms in Mendel's 1:2:1. Then the expected number of heterozygotes and mutant homozygotes can be calculated as Mendel's (1):2:1. The observed fraction of the expected number was then calculated. (Finally, the expected litter size was the product of the observed litter size for the 'normal group' and the observed fraction. For obvious reasons, this procedure was used only to estimate the magnitude of genotype loss; conventional chi-square tests were used to test the fit between observed and expected (1:2:1) genotype distributions. |  |  |  |  |  |  |  |  |  |  |

### Box 1. Genes that bias fertilization.

Additional information can be found in the Mouse Genome Database (informatics.jax.org).

#### A1cf

Apobec1 complementation factor, Chr 19, 26.6 cM. *A1cf* is expressed primarily in the nucleus where it encodes the RNA-binding subunit for APOBEC1 cytidine deaminase that edits specific bases, sometimes in coding sequences but more usually in 3’UTRs.^77, 210^ It must have additional functions since deficiency leads to early embryonic lethality,^48^ while APOBEC1-deficient mice are fully viable and fertile.^59, 60^ Partial deficiency increases TGCT risk in a parent-of-origin manner.^49^

#### Ago2

Argonaute RISC catalytic subunit 2, Chr 15, 33.9 cM. AGO2 is required for RNA-mediated gene silencing (RNAi) by the RNA-induced silencing complex (RISC).^79^ Guide RNAs (miRNAs and siRNAs) direct RISC to complementary RNAs that are targets for RISC-mediated gene silencing. AGO2 and PPP2CB (see below) promote mitotic chromosome segregation in the *Drosophila* and *C. elegans* germline.^98, 99, 211^ Interestingly, AGO3 piRNA component Aubergine enhances transmission distortion for the SD system in Drosophila,^212^ raising the possibility that AGO2 might have similar effects under appropriate circumstances. KRAS signaling controls AGO2 sorting into exosomes^213^ that transport RNAs, including tRNA fragments,^214^ for intercellular signaling. RNAs transferred to as well as produced in sperm could have significant effects on gamete functions.^215^ Selectivity in exosome targeting could lead to functional differences among haploid gametes. Loss of siRNA but not miRNA AGO2 activity leads to meiotic catastrophe in MI oocytes.^216-218^ In AGO2-deficient mice, miRNA levels are reduced substantially in oocytes,^219^ and in AGO2-deficient oocytes siRNA levels are reduced while retrotransposons and selected mRNA levels are increased.^220^ Homozygous deficient mice display embryonic lethality with various defects in embryonic and extraembryonic organs and tissues. Partial deficiency increases TGCT risk in a parent-of-origin manner.^49^

#### Apob

Apolipoprotein B, Chr 12, 3.53 cM. APOB is widely expressed where it transports lipids such as cholesterol. APOB is encoded as a single, long mRNA. The shorter apoB-48 protein is produced after RNA editing of the apoB-100 transcript at residue 2180 (CAA->UAA), resulting in the creation of a stop codon and early translation termination. Homozygous deficiency leads to embryonic lethality, with embryo loss by E9. Heterozygotes tend to have incomplete neural tube closure. Partial deficiency severely reduces fertility in males with sperm showing impaired motility and reduced ability to fertilize both *in vivo* and *in vitro*, arguing for a diploid rather than a haploid effect.^221^, ^222^

#### Apobec1

Apolipoprotein B mRNA editing enzyme, catalytic polypeptide 1, Chr 6, 57.7 cM. *Apobec1* encodes a cytidine deaminase that edits C to U (read as T in coding regions) primarily in 3’UTRs.^77^ A1CF is the RNA-binding protein that targets specific mRNA sites for editing.^48^ *Apobec1* interacts with the *Dnd1^Ter^* mutation to increase TGCT risk in a conventional manner in the male germline and in a transgenerational manner in the female germline.^59^

#### Dnd1

Dead-end microRNA-mediated repression inhibitor 1, Chr 18, 19.5 cM. *Dnd1* is an *A1cf-related* RNA-binding protein expressed in many tissues.^61^ DND1 controls access of particular miRNAs to their mRNA targets in human TGCTs.^80^ It is essential for germ cell differentiation^142^ and also acts like *A1cf* and *Ago2* in offspring to process inherited epigenetic changes from parents.^49^ DND1 works with NANOS2 in the CCR4-Not (CNOT) deadenylase complex to suppress specific RNAs.^223^ DND1 and NANOS2 load RNAs onto the CNOT complex for germ cell differentiation .^223^ The *Ter* mutation severely reduces fertility and is a potent modifier of TGCT susceptibility,^61, 224^ whereas the targeted deficiency results in biased fertilization.^45^

#### Ddx1

DEAD (Asp-Glu-Ala-Asp) box polypeptide 1, Chr 12, 6.4 cM. *Ddx1* is expressed primarily in the nucleus of the fetal and adult testis and ovary where it functions as an ATP-dependent RNA helicase to unwind RNA-RNA and RNA-DNA secondary structures for translation initiation, nuclear and mitochondrial splicing, ribosome and spliceosome assembly (including tRNAs), and pre-miRNA and polyA processing.^225, 226^ Reduced DDX1 activity promotes ovarian tumor growth.^226^ In Drosophila, *Ddx1* deficiency results in stress (starvation)-induced sterility in males and autophagy in egg chambers.^227^

#### L3P

No information except Marean et al.^76^

#### Lrp6

Low density lipoprotein receptor-related protein 6, Chr 6, 65.4 cM. *Lrp6* is expressed at E11.5 and after in female reproductive system and in fetal testes.^228^ *Lrp6* encodes a transmembrane cell surface protein involved in receptor-mediated endocytosis of lipoprotein and protein ligands. It can function alone or as a co-receptor with Frizzed for canonical Wnt/beta-catenin signaling. LRP6 and other members of the Hedgehog and WNT pathways are expressed in hESCs and testicular cancers.^229^ Partial embryonic lethality, growth retardation, and various vertebral and skeletal abnormalities are found in mutant homozygotes, and more subtle skeletal defects in mutant heterozygotes.

#### Ppp2cb

Serine/threonine protein phosphatase 2 catalytic subunit beta isoform, Chr 8, 20.6 cM. *Ppp2cb* encodes the 2A catalytic subunit of the PP2A heterodimer and is expressed in the female and male reproductive systems from E15 through adulthood. In particular, it is highly expressed in mature spermatozoa and MII oocytes where it localizes at centromeres in meiosis and spindle poles in mitosis.^230^ It is a negative regulator of the MAPK pathway and plays a role in DNA damage response, cell cycle control, apoptosis and mRNA splicing. Inhibition of PPP2CB releases meiotic arrest and enables meiotic progression.^230, 231^ Loss of PPP2CB in oocytes causes both failure of meiosis II exit and reduced fertility in females^230, 231^ and males.^232^ Related proteins play similar roles in meiotic control and chromosome segregation.^233-235^ PPP2CB deficiency affects sperm tails.^232, 236^ and chromosome segregation in females.^230^ Other reports find viable and fertile *Ppp2cb* mutant homozygotes, without obvious phenotype.^237^

#### Pum1

Pumilio1 RNA binding family member 1, Chr 4, 63.4 cM. *Pum1* is a widely expressed cytoplasmic protein found in the ovary and testis throughout fetal development and adulthood. It encodes an RNA binding protein that targets Pumilio Response Elements (PRE) in 3’UTRs to recruit both CCR4-NOT deadenylase, other deadenylases, and miRNAs such as miR221 and miR222 that together repress expression of genes such as p27 that maintain genome integrity.^238-240^ Interestingly, DND1 blocks access of miR221 to its p27 mRNA target,^238^ suggesting that DND1 acts competitively with PUM1 to control miRNA actions. In the testis, PUM1 acts as a post-transcriptional regulator of spermatogenesis by binding to the 3-UTR of mRNAs coding for regulators of TRP53^241^ and also suppresses caspase- and TRP53-apoptosis in germ cells.^17, 241^ It is involved in embryonic stem cell renewal by facilitating the transition from pluripotency to differentiation.^242^ PUM1 deficient males exhibit reduced weight of testes and seminiferous tubules, reduced number of sperm, and increased germ cell apoptosis and infertility.^17^ Interestingly, PUM1 contributes to the number of primordial ovarian follicles, meiosis, and reproductive competence in females.^243^ PUM1 contribute to antiviral response,^244^ suggesting that they might play a more general role in stress response to environmental and physiological conditions that often lead to transgenerationally inherited epigenetic changes.^245^

#### Vangl2

Vang-like planar cell polarity protein 2, Chr 1, 79.5 cM. VANGL2 is found in the plasma membrane and cytoskeleton where it provides directional signals to cilia.^246^ It is expressed in female and male reproductive systems from embryonic day E15 through adulthood. Both male homozygous mutants (*Lp/Lp*) and female heterozygous mutants (Lp/+) are sterile.^247^

#### Zic2

Zinc finger protein of the cerebellum 2, Chr 14, 66.0 cM, ZIC2 is expressed in the female and male reproductive system from embryonic day E15 through adulthood and represses transcription in the nucleus. ZIC2 promotes self-renewal of liver cancer stem cells by recruiting the nuclear remodeling factor (NURF) complex to activate the OCT4 pluripotency factor.^248^ It also enhances transcription to promote differentiation of embryonic stem cells in Drosophila^249^ and controls naïve versus primed pluripotency state.^200^ Deficiency results in neurulation defects and embryonic lethality in mice^76^ and holoprosencephaly in humans.^250^

